# The Linkage among Sea Transportation, Trade Liberalization and Industrial Development in the context of Carbon Dioxide Emissions: An Empirical Investigation from China

**DOI:** 10.1101/2020.07.13.200386

**Authors:** Salih Kalaycı, Cihan Özden

## Abstract

The major goal of this paper is to focus on the existing literature regarding the linkage between maritime, trade liberalization and industrial development in the context of CO_2_ by using econometrical model. In this context, it is attempted to reveal the effects of independent variables on CO_2_ (dependent variable) for China from 1980 to 2013 (annual data) by implementing Phillips-Perron (PP), Zivot-Andrews unit root tests, FMOLS, DOLS, CCR, ARDL and GMM methods. According to results of FMOLS, DOLS and CCR models there is a long-term stable relationship between sea transportation, trade liberalization, industrial development and carbon dioxide emissions which is proved empirically. Similarly, Short term ARDL estimation results reveal that the main determinants of CO_2_ in the short-run are changed in industrial development and maritime transport at 1% significance level. Table 6 summarizes the short-term ARDL results and the findings regarding the error correction model. According to Table 6, error correction model works in order to reach short-run adjustment. In the short term, approximately 78% of shocks in industrial development, maritime transport and trade liberalization are compensated within a period of time and the system is re-established in the long term. China produced half of the 1.2 million electric media used worldwide; the government directs its attention to the rehabilitation and reuse of all these lithium-ion batteries. Large-scale production of biofuels can still be several years away. Crude oil might be very difficult to promote alternative fuels on a national scale unless crude oil prices surge so high as to become unaffordable. Authorities underline: China will become the world’s number one economy. Now renewable energy will be more important, which should be encouraged to use by government on transportation so as to reduce the CO_2_ emissions. However, China can be leader excess oil use for transport if they want to dominate the economy worldwide.

## 1. Introduction

One of the most frequently discussed global issues in recent years has been environmental destruction in the context of global warming and climate change. The main cause of global warming is the very rapid increase in the rate of gases that cause the greenhouse effect in the atmosphere. The main gas causing the greenhouse effect through carbon dioxide (CO_2_) gas emitted to the atmosphere by the use of fossil fuels such as gasoline, coal and natural gas. Mass production and excessive consumption, which started through the industrial revolution, increased energy needs and this requirement was met largely from fossil fuels. For many years, fossil fuel energy demand has reached exponential growth, causing pollution on the environment. Thus, the countries that focused on the economic growth target also caused the CO_2_ emissions and global warming. The emergence of environmental pollution and industrialization has begun to be discussed in academic literature. One of the most important sources of this trend is the report “The Limits to Growth” prepared by the Roman Club in 1972. According to the report, “If the current growth of industrialization continues in terms of food production, consumption of natural resources and CO_2_ emissions, economic growth will damage the environment in our world in the next 100 years” [1].

Many studies have examined the dynamic relationships between transportation, CO_2_, industrial development and economic growth by taking into account the inverted U-shaped hypothesis as a base in the academic literature. However, the linkage among sea transportation, trade liberalization and industrial development in the context of CO_2_ will be new variables. China has witnessed tremendous economic growth, as well as rapid development in both the financial and labor sectors since last 20 years. Nowadays, China is the second largest economy in the world. Besides, economic growth was accompanied by increased fuel energy consumption and thus CO_2_ emissions.

In this study, empirical results proved the impact of sea transportation, trade liberalization and industrial development on carbon dioxide emissions for China from 1980 to 2013. In order to examine these dynamic relationships, the researchers applied many kinds of econometrical methods, such as Multivariate Regressions, the Johansen co-integration test, the ADF (Augmented Dickey–Fuller) unit root test, the VAR (Vector Autoregressive) model, impulse response analysis, variance decomposition analysis, Granger causality test, and panel data analysis in the methodology section of their articles. The researchers achieved different results for the validity of EKC (Environmental Kuznets Curve) relationships based on different samples, methodologies, and periods.

The major goal of this paper is to concentrate on the existing literature regarding the relationship between maritime, trade liberalization and industrial development in the context of CO_2_ by using econometrical model. In this context, it is attempted to reveal the effects of independent variables on CO_2_ (dependent variable) for China from 1980 to 2013. The main purpose of this analysis is to address the problem in terms of environmental economy by giving some suggestions. The study is structured as follows. Following this introduction, section 2 presents the theoretical background of relevant works and theorical framework of the article; section 3 deals with the analyzing and result of research and consequently section 4 concludes this article by giving some recommendation to reduce CO_2_ emissions.

## 2. Literature Review

Production forms are gradually divided into nations in today’s globalizing world. In this sense, local consumption in any country is incrementally performed throughout the world of supply chain. The transportation sector has developed through industrialization process and as a result carbon emissions has increased day by day especially in certain countries. Therefore, in this case, it imposed some restrictions in terms of trade and supply chain. Trade liberalization, transportation, industrial development and CO_2_ are widely discussed regarding the environmental consequences of trade in academic literature. Hassan and Nosheen [2] investigate the effects of trade openness energy usage and economic growth on carbon dioxide leakage in terms of EKC hypothesis for Pakistan by implementing yearly data beginning from 1980 to 2016. In order to reach the results they implement econometric models including ADF unit root test, Johnsen co-integration test to reveal the long-term relationship between variables and vector error correction. According to their findings, economic growth is positively connected with carbon dioxide leakage. Population growth and trade liberalization have a considerably negative effect on CO_2_ leakage during the short-term, whereas, in the long-term, the course is opposite. In addition, Granger causality test indicates that a bi-directional causality acts from energy consumption and economic growth to CO_2_ leakage. One-way causalities running from trade liberalization to CO_2_ emissions as well. Jebli and Belloumi [3] state that sea transportation, waste usage and combustible renewables significantly affect the carbon dioxide emissions, while any rise in sea transportion reduce the consumption of combustible renewables and waste usage. Besides, sea transportation is significantly correlated with carbon dioxide leakage, demonstrating that the Tunisian transportation is so contaminated due to the excessive non-renewable energy consumption. Consequently, sea transport is the main contributor to air pollution and lead to the increment of carbon dioxide leakage in the Tunisia. Nakayama, Zhu, Hirokawa, Irino and Yoshikawa-Inoue [4] assert that Rishiri Island is the northernmost site of the Japan where weather is measured, although it is not yet being incorporated by the WMO. Past works demonstrate that when the continental sea transport is performed especially in 90–150°E,40–60°N, it causes CO_2_ emissions. Zhu, Yuen, Ge and Li’s [5] the findings indicate that sea transport emissions in terms of trading system can motivate main actors to benefit from new technologies, provide more carbon efficient vessels in terms of green energy. Besides, the efficacy of sea transport emissions trading system is more obvious when bunker fuel prices are excessive. In this sense, the findings ascertain that bunker fuel prices have a larger impact on CO_2_ reduction than implementing stricter CO_2_ allowance allocation as well. Their work involves the formulation of sea transport emissions trading system policies and ensures suggestions in order to decrease the sea transport emissions to fulfill the shipping industry’s impetus by providing its environmental performance.

Katircioğlu [6] demonstrates that energy usage has positive while incoming tourists and economic growth have negative effect on carbon dioxide leakage in the long-run. It is further clarified that Singapore has an inverted U-shaped environmental Kuznets curve and regardless of the energy usage level, carbon dioxide leakage follow a declining trend in Singapore. Besides, the relationship among sustainability of the environment, energy efficiency and tourism industry development can be clarified by the tourism-induced environmental Kuznets curve hypothesis. According to which, in the beginning period of its trend, tourism sector lead to environmental pollution until the frontier point of country income is reached. In addition, as this point of income is reached however, it is expected to follow a downward point in the decline level.

Katircioglu, Katircioglu and Kilinc [7] express that industrial development, aggregate households, and therefore urban areas result in additional energy demand, which causes an expansion in carbon dioxide emissions. They empirically proved the urbanization-induced environmental Kuznets curve hypothesis and thus searched the long-run equilibrium linkage causality among CO_2_ emissions and urban development from energy usage and real income growth in the world. Katircioglu, Gokmenoglu and Eren [8] assert that tourism industry is relied majorly on infrastructure potentials including highways, airports, harbours, hotels and holiday village. However, it can be inevitable that tourism industry and its factors will considerably influence the environmental quality. Besides, abuse of the natural resources is one of the required operations in terms of attracting more individuals and ensuring a competitive tourism industry. Deforestation, abusing the raw materials and overutilizing the natural water are some of the compromises made to engender tourism industry by constructing hotels and other facilities. Thus, exploitation of the natural resources can cause undesirable environmental effects including excessive CO_2_ leakage, air pollution and erosion. Given its potential negative effects, connection among industrial development and environment quality has been examined in academic literature. In this sense, the findings of their work demonstrate that U-shaped is confirmed in the context of environmental Kuznets curve hypothesis for main tourist countries. Consequently, main tourist countries finely operate the tourism and urban development to control environmental pollution.

Katircioglu [9] indicates that tourism industry causes the rise of CO_2_ leakage for the Cyprus which is an economy known for its extreme demand of tourism activities. Furthermore, urbanism and entire population of the world have been involved in the research results for comparison aims as well. Koksal, Işik and Katircioğlu [10] state that developed economies concentrate more on light manufacturing industries including apparel, leather, wood, metal products and agribusiness. Relying on their phase of growth, the differentiation among these two economies allows them to differ their financial structure, which creates composition impact and Pollution Haven Hypothesis can clarify this fact. In this sense, liberal trade theories demonstrate that countries decide to manufacture products which they have a comparative advantage. Mehrara and Rezaei [11] examine the linkage among economic growth, CO_2_ emissions and trade openness through econometric models of unit root test, cointegration test and a panel data analysis from 1960 to 1996 for BRICS countries. In this context, data structure is tested to reveal the stationarity of series by implementing the ADF and PP unit root tests which demonstrates that the series are stationary at I(1). They reveal a cointegration linkage among CO_2_ emissions, economic growth and trade openness by using Kao panel cointegration test. The proof demonstrates that in the long-term trade openness has a positive considerably effect on CO_2_ emissions and impact of trade openness on emission.

Kuik and Gerlagh [12] find out the impact of trade openness on CO_2_ emissions by using econometrical models. They use quantitative estimates of CO_2_ emissions by taking into account the Kyoto Protocol and free trade through lowering the import tariff which is determined in the Uruguay Round of multilateral trade aggreement. In addition, lowering import tariff causes CO_2_ emissions and the costs of abating the trade-induced CO_2_ emission are relative to the welfare gains of free trade. In this context, analysis of the trade-induced CO_2_ emissions demonstrates distinct differences among emissions caused by lowering import tariff on energy products and by on non-energy products. It demonstrates distinct differences in leakage responses between developing countries as well. Managi, Hibiki and Tsurumi [13] point out that the effect of trade liberalization on environmental pollution by implementing the instrumental variables technique. They show that the effect is considerable in the long-run, after the dynamic adjustment process, although it is small in the short-run and trade is determined as an advantageous factor in terms of the environment in OECD economies. It has prejudicial impact, however, on CO_2_ leakage and sulfur dioxide SO_2_ in non-OECD economies, although it does lower biochemical oxygen demand (BOD) emissions in these countries. Consequently, trade liberalization affects CO_2_ leakage by way of the environmental regulation impact and capital influence.

Baek and Kim [14] reveal that trade volume to CO_2_ leakage and energy causality holds for the developed economies; changes in degree of trade liberalization lead to corresponding changes in the rates of growth in emissions and energy usage. In this sense, CO_2_ leakage and energy usage to trade volume is found to hold for the developing economies; any shocks in emissions and energy consumption influence the fluctuations in terms of trade liberalization. Shahbaz, Tiwari and Nasir [15] point out the impact of trade liberalization, economic growth and coal consumption on CO_2_ leakage by implementing time series analysis from 1965 to 2008 for South Africa. Autoregressive distributed lag (ARDL) bounds testing approach to cointegration has been implemented to analyze the long-term linkage between the variables while short-term dynamics have been examined by using ECM model. Their results verified the long-term linkage between the relevant variables. Findings indicate that an increase in GDP raises CO_2_ leakage, while financial development decreases it. Moreover, Coal consumption has remarkable effect to pollute the environment in South Africa. Trade liberalization factor has positive effect on environmental quality by decreasing the growth of energy pollutants. Finally, their findings confirm the existence of EKC.

Shen [16] demonstrate that factor endowment hypothesis is considered by several pollutants academic works, whilst there seems no proof of pollution haven hypothesis. Combining with all the estimated elasticities of scale and technique influences, composition impacts and trade intensity on CO_2_ leakage, found that raising trade volume has opposite effects on CO_2_ emissions because of different pollutants. Both SO2 and dust fall cause emissions, whilst trade openness decreases emissions. Oh and Bhuyan [17] examine the linkage among energy usage, GDP, trade liberalization, and carbon dioxide (CO_2_) leakage in Bangladesh from 1975 to 2013. They use ARDL bounds test to cointegration for establishing the existence of a long-term linkage. Conversely, the estimated coefficients for trade openness and GDP are negative both in short-term and long-term. They recommend that the government of Bangladesh should support the appropriate policy to use alternative energy facilities which would not emit much CO_2_.

Zandi and Haseeb [18] investigate the effect of trade openness on CO_2_ emissions. They implemented the panel data analysis for 105 developed and developing countries from the period of 1990 to 2017. The findings of FMOLS and DOLS verify that all variables are linked in the long-term. The findings of long-term coefficient verify that the trade openness has a positive impact on CO_2_ emissions and cause to increase environmental pollution. The findings of heterogeneous panel causality verify that there is a one way causal linkage among trade openness and environmental pollution where causality is running from trade openness to environmental pollution. Besides, they determine a two-way causal linkage of environmental pollution with renewable energy utilization in all chosen countries.

Ozatac, Gokmenoglu and Taspinar [19] examined the environmental Kuznets curve hypothesis for Turkey from the period of 1960 to 2013 by considering the energy usage, trade openness, financial development and CO_2_ emissions. The findings of the ARDL bound test and the error correction model through autoregressive distributed lag mechanism demonstrated long-term linkage between the relevant variables as well as evidence of the EKC hypothesis. They confirmed causal linkage between the variables and propose a “polluter pays” mechanism to maintain the awareness of a clean environment. Ali et al. [20] show that there is a statistically inconsequential correlation among CO_2_ emissions, GDP and industrial development of Pakistan. However, the result of Ali et al. [20]’s result is not consistent with this paper. Li et al. [21] discuss in detail the large and widening energy consumption, Air Quality Index and CO_2_ efficiency score gap among Chinese regions, through the lower CO_2_ and Air Quality Index efficiency scores mainly focused in the western cities. They recommend that China needs to pay more attention to the differing economic levels, social development, industrial structures, energy consumption, and R&D in the western regions and apply systematic solutions based on domestic meteorological and climatic conditions, economic and social development.

## 3. Methodology and Data Analysis

This manuscript examines the nexus between sea transportation, trade liberalization, industrial development and carbon dioxide emissions by implementing FMOLS, DOLS, CCR and ARDL model. Annual datas are obtained from UNCTAD’s [22] and Worldbank’s [23], [24], [25] official website. In this context, Phillips-Perron (PP) [26] and Zivot-Andrews [27] unit root tests are implemented in order to analyze whether variables are stationary. Afterwards, FMOLS, DOLS, CCR and ARDL method are used to reveal the co-integration connection among sea transportation, trade liberalization, industrial development and carbon dioxide emissions. In conclusion part, GMM test is undertaken to demonstrate the impact of sea transportation, trade liberalization and industrial development on carbon dioxide emissions for China from 1980 to 2013 yearly. In this manuscript, log-linear specifications of the variables are implemented to estimate the equation below:

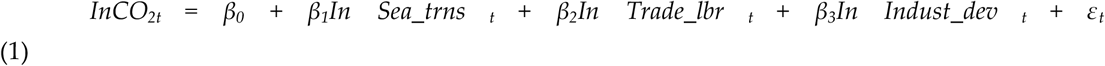

Where CO_2t_, Sea_trns_t_, Trade_lbr_t_, Indust_dev_t_ indicate carbon dioxide emissions, sea transportation, trade liberalization and industrial development respectively. *β*_*1*_, *β*_*2*_ and *β*_*3*_ provide the elasticity of the explanatory variables.

**Table 1.** Phillips-Perron (PP) Unit Root Test Results of China.

| Variables | PP (Fixed and trending)<br>(Level 0) | PP (Fixed and trending)<br>(First Difference) | Decision |
| --- | --- | --- | --- |
| ln Sea_trns | -2.9843<br>**(-3.9958)** | -6.5246*<br>**(-3.7245)** | I(1) |
| ln Trade_lbr | -1.4527<br>**(-5.2358)** | -4.1235**<br>**(-2.9152)** | I(1) |
| ln Indust_dev | -1.9751<br>**(-3.9614)** | -5.7286*<br>**(-3.2724)** | I(1) |
| ln CO<sub>2</sub> | -3.1262<br>**(-3.8261)** | -6.5192*<br>**(-3.4786)** | I(1) |
| Note: - * and ** Expressions define the unit root test results of the variables used in the estimation process, 1% and 5% significance levels, respectively. |  |  |  |

**Table 2.**
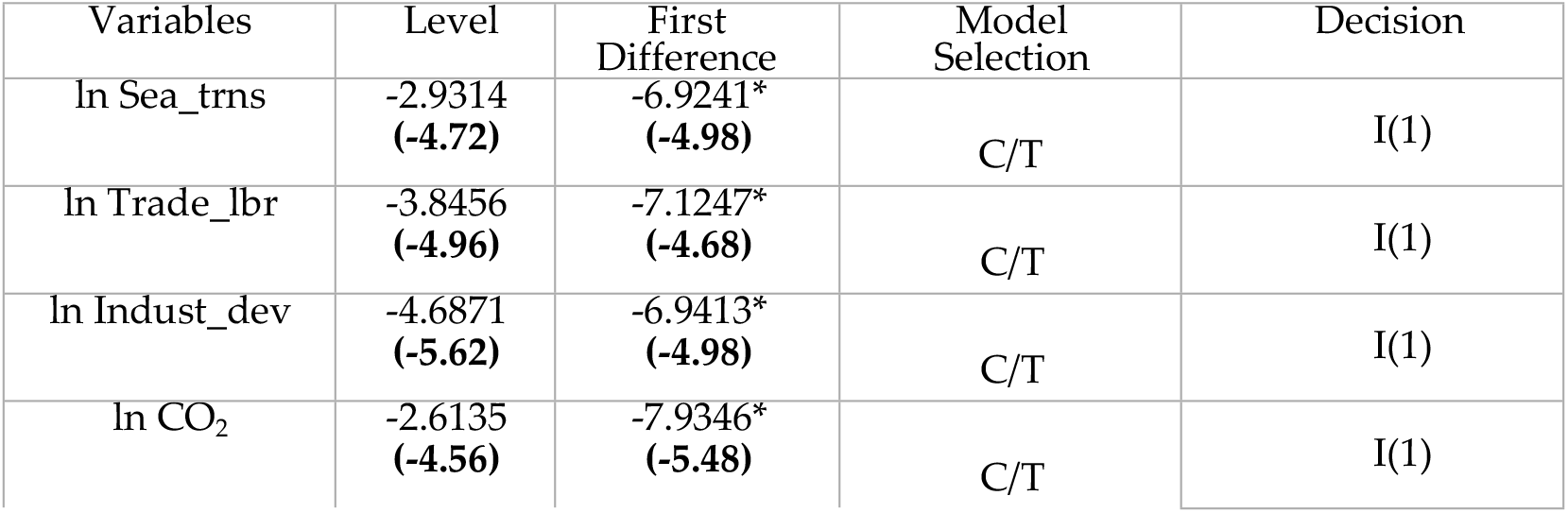
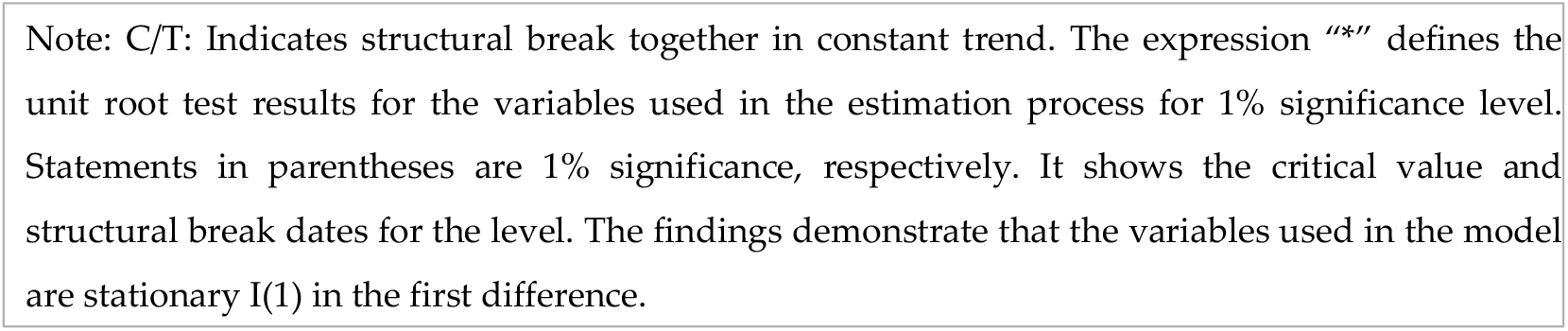
Zivot-Andrews Unit Root Test Results of China.

| Variables | Level | First Difference | Model Selection | Decision |
| --- | --- | --- | --- | --- |
| ln Sea_trns | -2.9314<br>**(-4.72)** | -6.9241*<br>**(-4.98)** | C/T | I(1) |
| ln Trade_lbr | -3.8456<br>**(-4.96)** | -7.1247*<br>**(-4.68)** | C/T | I(1) |
| ln Indust_dev | -4.6871<br>**(-5.62)** | -6.9413*<br>**(-4.98)** | C/T | I(1) |
| ln CO<sub>2</sub> | -2.6135<br>**(-4.56)** | -7.9346*<br>**(-5.48)** | C/T | I(1) |

Note: C/T: Indicates structural break together in constant trend. The expression “\*” defines the unit root test results for the variables used in the estimation process for 1% significance level. Statements in parentheses are 1% significance, respectively. It shows the critical value and structural break dates for the level. The findings demonstrate that the variables used in the model are stationary I(1) in the first difference.

Phillips-Perron (PP) [26] test is undertaken in order to reveal the structure of series in terms of stationarity. In this sense, according to Phillips-Perron [26] test results all series including sea transportation, trade liberalization, industrial development and carbon dioxide emissions are not stationary. For this reason, first differences of all series are taken to comprehend the structure of series in terms of stationarity. According to results of Phillips-Perron (PP) [26] test, series become stationary after taking first differences of variables. Thus, FMOLS, DOLS, CCR test can be implemented. However, performing traditional Phillips-Perron (PP) [26] unit root test without considering these structural breaks in the model may give false results and lose reliability in terms of estimation [26]. Therefore, Zivot and Andrews [27] develop a series of tests to consider endogenous structural changes. The test allows the evaluation of the presence of a unit root against the alternative of a stationary process through a structural change both in level or trend. Thus, the ZA test investigates the possibility of the existence of a segmented trend. ZA test tries to determine the structural break and treat it as endogenous in the selected sample. The following equation regarding ZA test is indicated below;

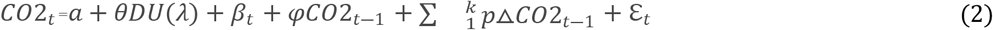

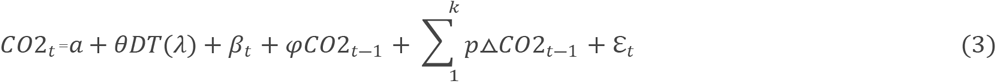

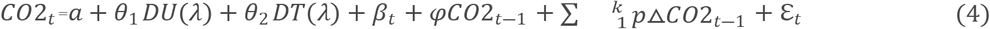

Consequently, the model for CO_2_ emissions in China would be as follows:

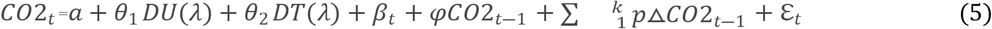

CO_2_ is the logarithm of carbon dioxide emissions expressed in levels, α is a constant, DU (λ) is a dummy variable that takes the value as 1 from the series in which the structural change is considered to have the value of 0 in previous years, the variable of t represents the time, CO_2_ −1 is carbon dioxide emissions lagging one period. DT (λ) = t − Tλ if t > Tλ and 0 if this is not the case. The next term is the sum of the change in the variable of interest for periods t − j through k; The regressors of this term are added to eliminate the possible dependence on the limit distribution used in statistical tests, caused by the temporal dependence of the distributions. Finally, ε is the error term.

According to results of FMOLS, DOLS and CCR models there is a long-term stable relationship between sea transportation, trade liberalization, industrial development and carbon dioxide emissions which is demonstrated empirically. The p-value of sea transportation, trade liberalization and industrial development are less than 0.05 at Table 3. Thus, there independent variables affect the dependent variable (CO_2_). Considering the coefficients of the FMOLS model; 1 percent increase in industrial development increases CO_2_ by 0.1678 percent, a 1 percent increase in trade liberalization increases real CO_2_ by 0.2862 percent, a 1 percent increase in maritime transport increases real CO_2_ by 0.3784 percent. According to coefficients of the DOLS model; a 1 percent increase in industrial development increases CO_2_ by 0.2156 percent, a 1 percent increase in trade liberalization increases real CO_2_ by 0.3374 percent, and a 1 percent increase in maritime transport increases real CO_2_ by 0.2979 percent. Finally, considering the coefficients of the CCR model; a 1 percent increase in industrial development increases CO_2_ by 0.1879 percent, a 1 percent increase in trade liberalization increases real CO_2_ by 0.3247 percent, a 1 percent increase in maritime transport increases real CO_2_ by 0.3216 percent.

**Table 3.** Cointegration Estimation Results (FMOLS, DOLS and CCR Models of China).

| <u>Variables</u> |  |  |  |  |  |  |  |  |  |
| --- | --- | --- | --- | --- | --- | --- | --- | --- | --- |
| <u>Dependent</u><br>CO<sub>2</sub> | Fully modified least square<br>(FMOLS) |  |  | Dynamic least square (DOLS) |  |  | Canonical<br>co-integrating<br>regression |  |  |
| <u>Independent</u><br><u>Variables</u> | <u>T-stat</u> | <u>P-val</u> | <u>Coef</u> | <u>T-stat</u> | <u>P-val</u> | <u>Coef</u> | <u>T-stat</u> | <u>P-val</u> | <u>Coef</u> |
| Sea_trns | -0.693462* | 0.0257 | 0.3784 | -0.617731* | 0.0482 | 0.2979 | -0.786156* | 0.0317 | 0.3216 |
| Trade_lbr | -1.842746* | 0.0486 | 0.2862 | -2.678134* | 0.0179 | 0.3374 | -2.124734* | 0.0454 | 0.3247 |
| Indust_dev | -1.23575* | 0.0368 | 0.1678 | -1.678965* | 0.0456 | 0.2156 | -1.896543* | 0.0246 | 0.1879 |
| C | 2.962458* | 0.0013 | 1.9321 | 3.124237* | 0.0306 | 2.6479 | 3.118965* | 0.0067 | 1.7895 |

ARDL method is implemented out of FMOLS, DOLS and CCR analysis whether there is a long-term causality relationship between sea transportation, trade liberalization, industrial development and carbon dioxide emissions. Table 5 presents the results obtained with the ARDL method below. Results of ARDL model are consistent with FMOLS, DOLS and CCR analysis.

FMOLS, DOLS and CCR co-integration analysis depend on the condition that the series implemented, such as traditional co-integration method which is required stationary series. In addition, having the possibility to interpret the derived coefficients provide course of process in terms of CO_2_ emissions through considering independent variables including sea transportation, trade liberalization and industrial development. The ARDL equation is indicated as econometric symbols, where the determinants of long-term economic growth are investigated in equation (6) below:

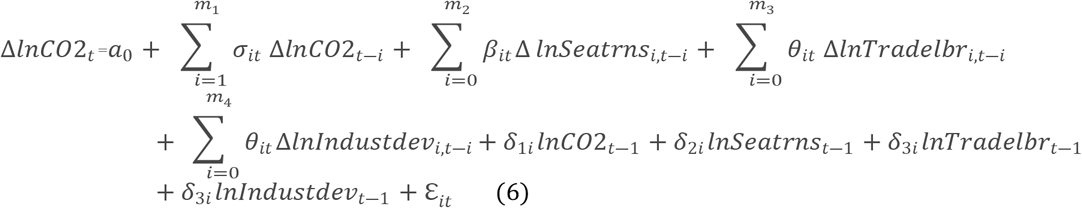

The long-run relationship between carbon dioxide emissions *CO*2_*t*_ sea transportation *Seatrns*_*t*_ trade liberalization *Tradelbr*_*t*_ and industrial development *Industdev*_*t*_ is investigated through f bounds test which is considered the zero hypothesis.

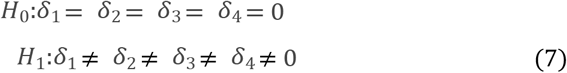

Findings from FMOLS, DOLS and CCR models indicate that maritime transport, trade liberalization and industrial development are the determinants of long-term carbon emissions, just as in the results of the ARDL model. It is also noteworthy that the findings obtained from FMOLS, DOLS and CCR models, which are described as new co-integration techniques and allowed the separation of short and long-term relationships, consistent with the long-term results obtained from the ARDL model in Table 5. In this context, trade relations should be increased by policy-makers through improving their maritime transport infrastructures and further accelerate their industrial growth via research and development.

Considering the ARDL F-bound test, long-term ARDL estimates were made, respectively, through the revealing long-term co-integration by empirical model which is used among the variables. Long term ARDL forecast results are given in Table 4. Long-term ARDL forecast results reveal that the main determinants of CO_2_ changes in sea transportation, trade liberalization, industrial development.

**Table 4.**
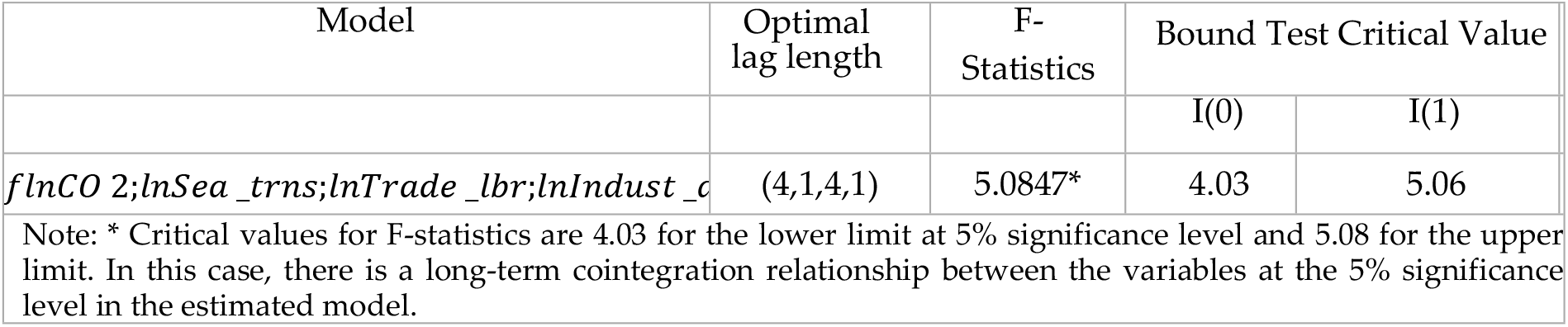
ARDL Bound Test Results of China.

According to the long-run ARDL results summarized in Table 5; If sea transport increases by 1 percent, CO_2_ increases by 0.3017%. A 1 percent increase in industrial development increases CO_2_ by 0.2972%. Unlike other econometric models (FMOLS, DOLS and CCR), trade liberalization has no statistically significant effect on CO_2_. Long-run ARDL results in terms of the relationships between CO_2_ and key economic determinants are similar with short-term ARDL test findings. Short term ARDL estimation results reveal that the main determinants of CO_2_ in the short-run are changed in industrial development and maritime transport at a 1% significance level. Table 6 summarizes the short-term ARDL results and the findings regarding the error correction model. According to Table 6, error correction model works in order to reach short-run adjustment. In the short term, approximately 78% of shocks in industrial development, maritime transport and trade liberalization are compensated within a period of time and the system is re-established in the long term.

**Table 5.** Long-Term ARDL Estimation Results of China.

| Dependent Variable: $\ln CO_2$ | | | | |
| --- | --- | --- | --- | --- |
| Variables | Coefficient | Standard error | t-Statistics | P-Val |
| Long-run Results |  |  |  |  |
| $\ln Sea\_trns$ | 0.3017 | 0.0372 | 3.9562 | 0.0021* |
| $\ln Trade\_lbr$ | -0.0756 | 0.2178 | -0.3681 | 0.5987 |
| $\ln Indust\_dev$ | 0.2972 | 0.0728 | 4.9852 | 0.0012* |
| Constant | 3.1673 | 0.7235 | 2.9641 | 0.0035* |
| Trend | 0.0059 | 0.0032 | 2.9246 | 0.0031* |
| Note: * expression defines the unit root test results of variables used in the estimation process at 1% significance level. |  |  |  |  |

**Table 6.** Short Term ARDL Results and Error Correction Model of China.

| Dependent Variable: lnCO<sub>2</sub> |  |  |  |  |
| --- | --- | --- | --- | --- |
| Variables | Coefficient | Standard error | t-Statistics | P-Val |
| Short-run Results |  |  |  |  |
| D (lnCO<sub>2</sub> (-1)) | 0.2144 | 0.0926 | 1.2856 | 0.2124 |
| D (lnCO<sub>2</sub> (-2)) | 0.1785 | 0.0868 | 1.9825 | 0.0356** |
| D (lnCO<sub>2</sub> (-3)) | 0.3286 | 0.1162 | 2.3175 | 0.0437** |
| D (lnIndust_dev) | 0.5267 | 0.0679 | 5.9245 | 0.0089* |
| D (lnSea_trns) | 0.1877 | 0.0518 | 2.8954 | 0.0000* |
| D (lnSea_trns (-1)) | 0.0042 | 0.0784 | 0.0317 | 0.8965 |
| D (lnSea_trns (-2)) | -0.2169 | 0.0829 | -2.2886 | 0.0467** |
| D (lnSea_trns (-3)) | -0.1657 | 0.0736 | -3.1627 | 0.0491** |
| D (lnTrade_lbr) | -0.2878 | 0.1973 | -1.9348 | 0.0965*** |
| Constant | 3.1674 | 0.4817 | 3.9523 | 0.0089* |
| Trend | 0.0089 | 0.0028 | 5.0167 | 0.0037* |
| CointEq(-1) | -0.7319 | 0.2147 | -3.8176 | 0.0026* |
| Note: *, ** and *** expressions indicate 1%, 5% and 10% significance levels, respectively. |  |  |  |  |

In final part of methodology, the generalized moments method (GMM) is implemented in order to reveal the linear relationship between variables. In econometrics, the generalized moments (GMM) is a general used analysis in order to estimate the parameters in terms of a statistical approach. In this context, generalized moments method (GMM) is a form of statistical model which pairs up the macroeconomic data with the information of population moment conditions to forecast the unknown components of the economic analysis. After obtaining these components, it should be investigated at the probability values to make inferences regarding the basic questions.

(GMM) the equation of linear relationship among relevant variables; Instrumental variables are;

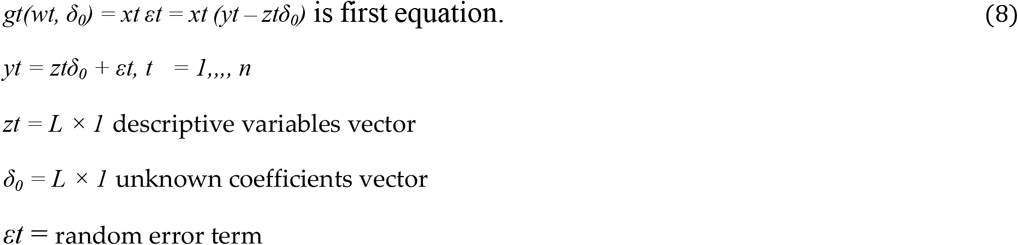

Instrumental variables are;

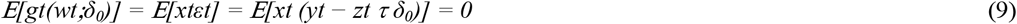

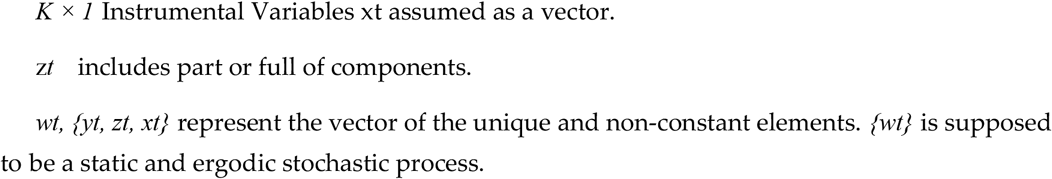

Time series may have self-relationships, and it can be said that this can lead to false results and endogeneity problems. GMM technique has been applied to minimize the internality problem. Different TSLS (Two-Stage Least Squares), White and HAC, were implemented by applying different GMM methods by choosing TSLS method. The analysis results of GMM are indicated below (Table 7).

**Table 7.** GMM Root Mean Square Error Results of China.

| GMM Methods Errors Comparison | Root Mean Square Error(RMSE) |
| --- | --- |
| GMM/TSLS | **4.96** |
| GMM/White | 6.72 |
| GMM/HAC | 5.32 |

Tablo 7, the lowest root mean square error is found in the GMM-TSLS methods. Therefore, GMM-TSLS method is selected for the analysis. According to GMMTSLS analysis (Table 8), there is no validation problem since the t-statistic value is more than 0.05. AR (1) is significant, and AR (2) is meaningless. The correlation among the variables of time series and the variables leading to or behind by a constant amount of time in terms of these parameters has been confirmed clearly. Furthermore, when considering the generalized moments method - TSLS (TwoStage Least Squares) model, there is no autocorrelation problem since the Durbin Watson value (1.76 - see Table 5.) is close to 2.

**Table 8.** GMM (Generalized Method of Moments) Test Results of China.

| Dependent Variable: CO<sub>2</sub>, Applied Analysis: GMM, Sample Size: 1980 – 2013, Observations: 34 |  |  |  |  |
| --- | --- | --- | --- | --- |
| Estimation Weighting Matrix: TSLS Two Stage Least Squares |  |  |  |  |
| Standard Errors & Covariance Computed Using Estimation Weighting, Matrix: C D(GDP) D(Ext_debt) D(Energy) |  |  |  |  |
| Variables | Coefficient | Std. Error | t-Statistic | Prob. |
| C | -8968554. | 29243784. | 4.235713 | 0.0028 |
| Sea_trns | 2.124562 | 0.317263 | 5.263267*** | **0.0092** |
| Trade_lbr | 0.673212 | 0.335861 | 3.261862*** | 0.0062 |
| Indust_dev | -0.371519 | 0.076314 | -3.547268*** | 0.0042 |
| AR(1) | -0.767874 | 0.296781 | -2.975379 | 0.0064 |
| AR(2) | -0.619623 | 0.213673 | -1.435296 | 0.3267 |
| R-squared | 0.819672 |  | Mean dependent var | 2.163681 |
| Adjusted R-squared | 0.786532 |  | S.D. dependent var | 1.471676 |
| S.E. of regression | 0.163578 |  | Sum squared resid | 13.1628 |
| Durbin-Watson stat | 1.761825 |  | J-statistic | 0.251231 |

Instrumental variables are added to the analysis as well. For the periodic intervals, three dummy variables are implemented as well. The first dummy variable is GDP, which indicates China’s total economic growth from 1980 to 2013. Another dummy variable indicates the overall external debt of China for the years 1980-2013. Energy consumption for China is used as the last dummy variable. In the generalized moments method, sea transportation, trade liberalization, industrial development influence CO_2_ in China which is considerably significant results.

## 4. Conclusion

Transforming the transport industry to run on renewable energy is vital for a more sustainable society and many countries are concerned with their carbon footprint. The clean energy revolution is also sweeping through the Chinese transport sector. In 2017 China produced half of the 1.2 million electric media used as worldwide; the government directs its attention to the rehabilitation and reuse of all these lithium-ion batteries. Large-scale production of biofuels can still be several years away. Crude oil might be very difficult to promote alternative fuels on a national scale unless crude oil prices surge so high as to become unaffordable. Developing battery science, solar panels and electricity are obtained from the sun rays. This obtained energy is also included in the systems to assist the ship’s shipbuilding auxiliary power or ship’s daily use needs. Some certain industries are using the solar panels for production. First looked at the solar panel due to the attitude and the cost of initial setup, it is necessary to actively wait for the future in the maritime industry. Therefore, first step reuse of waste energy CO_2_ emissions can be reduced. Thus, since the amount of fuel consumed by the ship decreases, the amount of CO_2_ emissions decreases. Moreover, methods to reduce CO_2_ emissions include reducing fuel costs and using alternative fuels with low or zero carbon content. Therefore, carbon emissions will be greatly reduced with burning low emission’s fuel in diesel engines.

Power management can be achieved by ensuring that diesel machines on ships operate at optimum powers. When machines are operated with excessive continuous overload, they can also be increased carbon dioxide emissions. Common Rail (Powerline) working principle includes the minimum fuel for each load sent to the cylinder regardless of the load the machine is running. It is possible to achieve 1% energy efficiency with this system. It is used not only to reduce CO_2_ emissions, also to reduce SO2 emissions. It is possible to achieve 1% energy efficiency with this system. Four types of engines are used in the manufacturing industry in the world, and these are equivalent engines IE1, IE2, IE3 and IE4, IE4 motors are the least energy consuming and most efficient motors in the world. In order to make the use of IE4 engines more common, efforts to reduce the cost of such machines should be given importance. However, roller bearing is usually found in automated machines such as CNC, and their usage rate should be increased with R&D supports by government. Intelligent start-stop systems are widely used today. Logically, the software is thrown in batteries electronic circuit, so that parameters are protected without the need for high energy. Thus, the machine start-up setup time is eliminated, and the machine can be used for longer production. This system should be applied in all machines with PLC panels and even adding start-stop batteries to the low-tech machines together with the panel will provide CO_2_ emissions, energy consumption and high profitability. Uninsulated and high-volume environments are cooled by fans in enterprises, resulting in high energy consumption and CO_2_ emissions. As an alternative system, the environment cooling system that works by rotating cold water in the company is much cheaper and environmentally friendly.

Findings from FMOLS, DOLS and CCR models indicate that maritime transport, trade liberalization and industrial development are the determinants of long-term carbon emissions, just as in the results of the ARDL model. It is also noteworthy that the findings obtained from FMOLS, DOLS and CCR models, which are described as new co-integration techniques and allowed the separation of short and long-term relationships, consistent with the long-term results obtained from the ARDL model in Table 5. According to the long-run ARDL, if sea transport increases by 1 percent, CO_2_ increases by 0.3017%. A 1 percent increase in industrial development increases CO_2_ by 0.2972%. Unlike other econometric models (FMOLS, DOLS and CCR), trade liberalization has no statistically significant effect on CO_2_. Long-run ARDL results in terms of the relationships between CO_2_ and key economic determinants are similar with short-term ARDL test findings. Short term ARDL estimation results reveal that the main determinants of CO_2_ in the short-run are changed in industrial development and maritime transport at a 1% significance level.

The impact of Covid-19 outbreaks with a prolonged shutdown of business operation negative effected China’s economy. Especially, manufacturing, transportation, trade and communication services and other top industries are suffered dramatically. Despite of Coronavirus pandemic, manufacturing in China is showing an upward trend, according to the statistics for March and April of 2020. China suffered great commercial damage due to the Covid-19. Now, China has increased its sales prices in all medical products and then other China’s origin products followed these increase prices which will not drop again due to Covid-19 even if China covers the loss. Authorities underline that China will become the world’s number one economy. As of now renewable energy will be more important, which should be encouraged to use by government on transportation (sea-railway-road) so as to reduce the CO_2_ emissions. However, China can be leader for excess oil use for transport if they want to dominate the economy worldwide.

